# Persistent transcriptomic changes following repeated exposure to wood smoke in non-human primate airway epithelial cells

**DOI:** 10.1101/2025.11.14.688533

**Authors:** Evan Holmes, Anthony P Brown, Sweeney P Elston, Christopher M Royer, Hong Ji

**Affiliations:** California National Primate Research Center, University of California Davis, Davis, CA, USA; Department of Anatomy, Physiology and Cell biology, School of Veterinary Medicine, University of California Davis, Davis, CA, USA

**Keywords:** Wood smoke exposure, Airway epithelial remodeling, Gene expression, DNA methylation, Ferroptosis, Epithelial barrier integrity

## Abstract

Exposure to wildfire smoke is a significant public health concern with the respiratory system heavily impacted. Wildfire events are increasing in number and severity, and inhalation of wildfire smoke can lead to airway inflammation, oxidative stress-induced lung injury, impaired mucociliary clearance, and exacerbation of respiratory conditions such as asthma and chronic obstructive pulmonary disease (COPD). Despite these well-documented health risks, the molecular mechanisms underlying the respiratory effects of wildfire smoke exposure and recovery remain incompletely understood. In this study, we utilized transcriptomic and epigenomic analyses to investigate how exposure to wood smoke (WS) influences the expression patterns of gene networks in differentiated tracheobronchial epithelial cells. Our analysis identified exposure-induced differentially methylated regions, differentially expressed genes and gene networks that are implicated in lung disease, including dysregulation of immune responses, oxidative stress and cell death, compromise of epithelial barrier integrity and function, and epigenetic remodeling. Strikingly, significant transcriptomic changes were still detected one week after exposure cessation, enriched in pathways involved in inflammation, wound healing and tissue repair. Despite these transcriptomic and epigenomic perturbations, histology staining revealed no significant changes in epithelial tissue morphology following WS exposure. However, we found a significant number of pulmonary disease-associated genes and pathways whose transcription were affected by WS exposure. Our study enhances understanding of the molecular basis of wildfire smoke-induced respiratory effects and highlights the potential for WS to leave a lasting imprint on airway epithelium, with important implications for respiratory health in exposed populations.

## Introduction

Wildfire events have become more frequent and severe in recent years, increasing concerns for public health (McClure & Jaffe, 2018). In California, the average annual area burned by wildfires has increased roughly 3.5-fold in the past decade, which resulted in a record 4.2 million acres burned in 2020 (OEHHA, 2022). Some regions of the state experienced over 90 smoke-filled days from this extreme fire season. Wildfire smoke is a complex mixture of particulate matter and toxic gases that currently contributes nearly half of the fine particulate pollution that is 2.5 micrometers or smaller in aerodynamic diameter (PM2.5) in the Pacific Northwest (Ford et al., 2018). Exposure to heavy wildfire smoke has been associated with substantial acute health burdens. Epidemiologic studies link smoke episodes to spikes in respiratory hospitalizations and to exacerbations of chronic respiratory diseases such as asthma and chronic obstructive pulmonary disease (COPD) (Aguilera et al., 2021, Reid et al., 2016, Roscioli et al., 2018, Wang et al., 2025). Although chronic effects remain incompletely characterized, there is growing concern that recurrent smoke exposures may contribute to longer-term health risks (Juarez et al., 2025, Kim et al., 2017, Orr et al., 2020), potentially accelerate the development or progression of chronic respiratory conditions.

Current understanding of the biological impacts of wildfire smoke exposure points to the respiratory epithelium as a primary target of injury. Abundant fine particulate matter in wildfire smoke is capable of depositing deep in the airways, resulting in acute inflammation and oxidative stress in the airway mucosa (Valavanidis et al., 2008). Airway epithelial cells exposed to wildfire smoke or wood smoke (WS) *in vitro* and *in vivo* show increased production of proinflammatory cytokines, induction of oxidative damage, and cytotoxic effects that disrupt the protective epithelial barrier (Heibati et al., 2025). These processes can lead to immune cell recruitment and broad activation of innate immune pathways in the lungs, which contributes to local tissue damage and systemic inflammation. Additionally, emerging evidence suggests that wildfire smoke can induce epigenetic remodeling in exposed cells. Environmental particulate exposures are known to cause epigenetic alterations such as changes in DNA methylation, which may influence gene regulation and disease susceptibility (Heßelbach et al., 2017). However, the timing, persistence, and mechanisms of such epigenetic changes in the context of wildfire smoke are not well understood. For example, it is unclear how long smoke-induced gene expression changes persist after an exposure, whether repeated exposures lead to cumulative or adaptive molecular responses, and what links exist between inhaled smoke and stable epigenetic modifications in airway cells.

These gaps in knowledge highlight the need for integrative molecular studies of wildfire smoke exposure. Nonhuman primates provide a valuable experimental model for investigating respiratory smoke exposure. Pulmonary anatomy, physiology, and immune responses of nonhuman primates closely parallel those of humans, permitting translational insights under controlled exposure conditions. Our prior research in rhesus macaques has demonstrated that ambient wildfire smoke exposure can have lasting biological effects when exposure occurs during early life. We found that infant macaques exposed to ambient severe wildfire smoke developed persistent methylation alterations in their airways that were still detectable years later (Brown et al., 2021). This underscores the potential for wildfire smoke to imprint long-term changes, especially during vulnerable developmental windows. However, adult lungs may respond differently than developing lungs, and it remains unclear whether repeated smoke inhalation in mature individuals elicits only transient perturbations or can also instigate lasting changes in gene regulation. Given the complexity of wildfire smoke’s effects, an integrative approach examining both gene expression and epigenetic alterations is warranted. In this study, we exposed differentiated adult rhesus macaque airway epithelial cells directly to diluted whole WS as a surrogate for wildfire. We characterized gene expression dynamics to delineate the temporal pattern of transcriptional responses. In parallel, we evaluated genome-wide DNA methylation in repeatedly exposed cells to determine whether smoke exposure induces epigenetic alterations that might underlie or exist beyond the measured transcriptional changes.

## Results

### Exposure Level Analysis

Tracheobronchial epithelial cells isolated from adult rhesus monkeys (ages>3 yrs) were cultured and differentiated at air-liquid interface (see Methods) and exposed to WS six hours per day for 1 day, 5 days and 5 days followed by 7 days of recovery (Figure 1). Gravimetric analysis of filter sampling during the six-hour exposures indicated chamber total particulate levels were 82.7, 242.8, 157.2, 168.1, and 182.5 μg/m^3^ for exposure days 1 to 5, respectively. The average of 166.7 μg/m^3^ for the five-day exposure is consistent with an “unhealthy” air quality index rating as indicated by National Ambient Air Quality Standards from the Environmental Protection Agency. The single day exposure group was included on day 2 (242.8 μg/m^3^) of the 5-day period and was consistent with a “very unhealthy” air quality index. Chamber carbon monoxide levels averaged 2.7, 9.1, 11.5, 11.2, and 4.9ppm for exposure days 1 to 5, respectively. Particle counts were 489.4/ml (SD 41.0; 0.3-0.5 μm), 41.8/ml (SD 33.7; 0.5-1 μm), 1.4/ml (SD 0.87; 1-5 μm), 0.04/ml (SD 0.02; 5-10 μm), and 0 for 10-25μm and >25 μm particle size bins.

**Figure 1:**
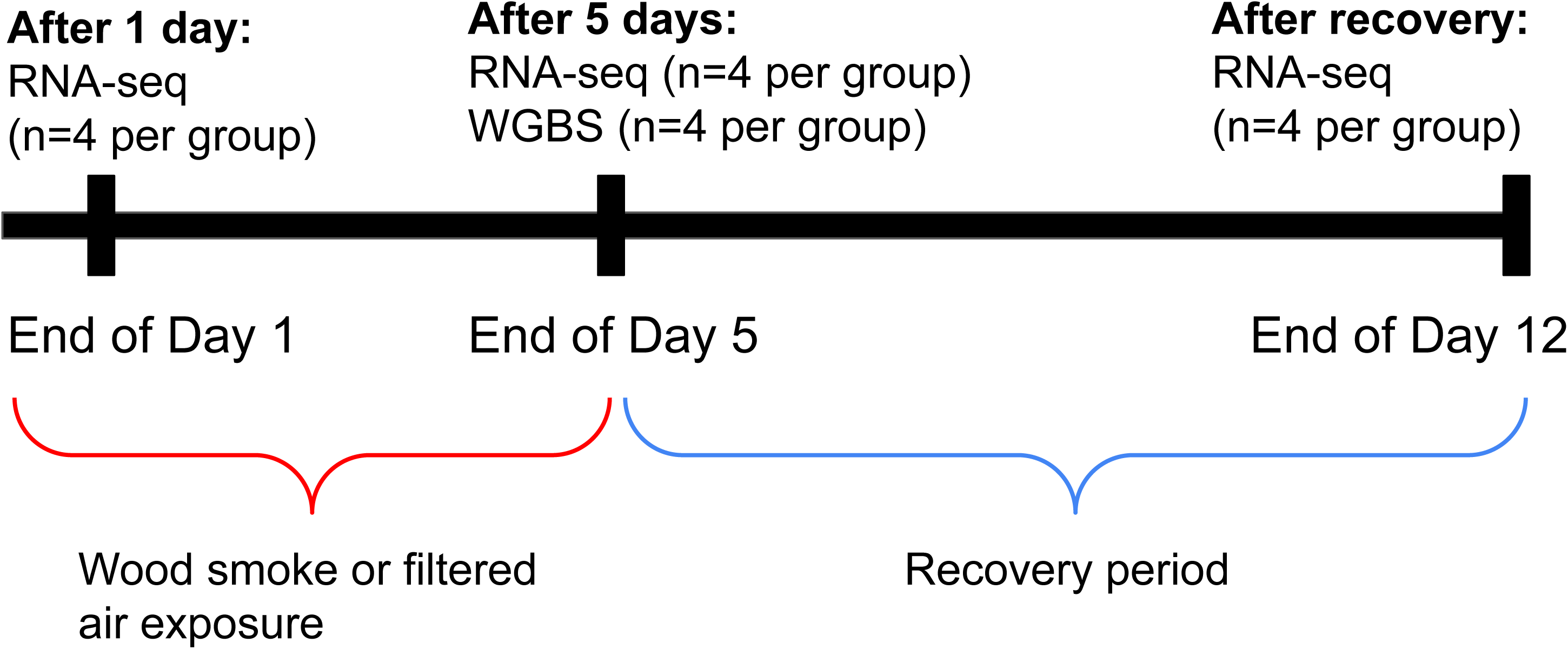
Overview of study design. Differentiated rhesus macaque airway epithelial cells were exposed to WS for six hours per day for 1 day, 5 consecutive days, or allowed to recover for 7 days post 5-day exposure. WGBS and RNA-seq were performed at the 5-day timepoint and all treatment timepoints, respectively.

### Wood Smoke Exposure Induces Significant Changes in Gene Expression

To examine the effects of WS exposure on gene expression, we conducted RNA sequencing analysis on these exposed airway epithelial cells (Figure 1). At the 1-day timepoint, 39 DEGs (|FC| ≥ 1.2 & adjusted p<0.05, Figure 2A and Supplementary Table 1A) were identified, reflecting early transcriptional changes immediately following exposure. Ingenuity pathway analysis (IPA) analysis revealed that these genes were significantly enriched for pathways involved in aryl hydrocarbon receptor signaling, inflammation, and oxidative stress (Supplementary Table 2A). At the 5-day timepoint, we identified 615 DEGs (|FC| ≥ 1.2 & adjusted p<0.05, Figure 2B and Supplementary Table 1B), representing transcriptional changes in response to repeated WS exposure. These genes were enriched for pathways involved in aryl hydrocarbon receptor signaling, cell division regulation, and ferroptosis signaling (Supplementary Table 2B). Twelve genes overlapped in response to exposure for 1 and 5 days (Figure 2D), and Aryl Hydrocarbon Receptor (AhR) Signaling was enriched among both sets of DEGs.

**Figure 2:**
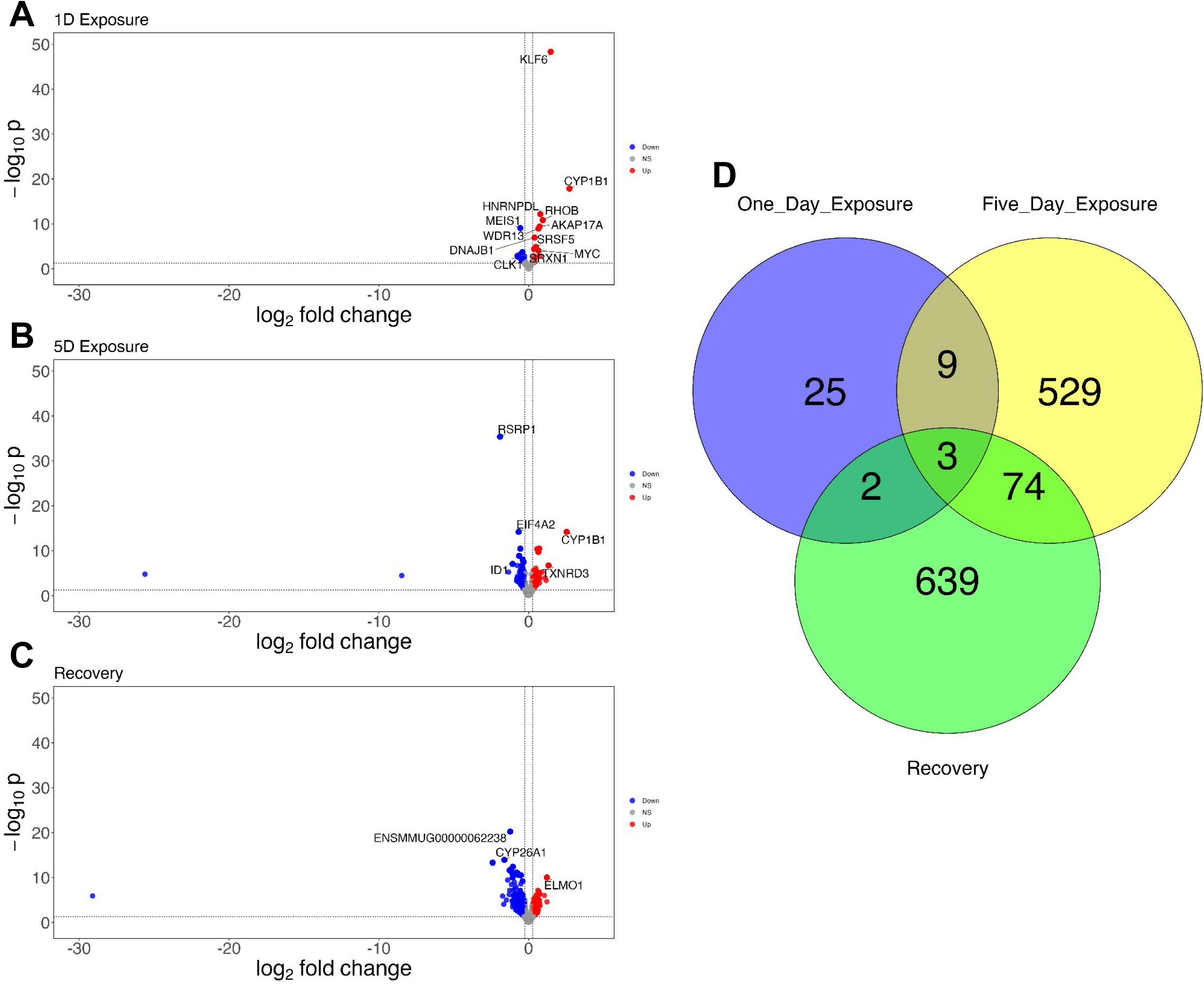
WS smoke induces persistent changes in gene expression. A) Volcano plot showing differential expression following 1 day of exposure to WS. B) Volcano plot showing differential expression following 5 days of exposure to WS. C) Volcano plot showing differential expression following a one-week recovery period from 5 days of repeated exposure to WS.

At the recovery time point, 718 DEGs were identified (Figure 2C and Supplementary Table 1C). Interestingly, enrichment of these genes indicated the downregulation of pathways involved in pulmonary healing signaling, TGF-β signaling, and regulation of the EMT by growth factors (Supplementary Table 2C). Notably, 77 DEGs overlapped between the 5-day and recovery time points (Figure 2D) and 27 featured changes in the same direction across exposure groups. IPA of these overlapping DEGs revealed significant enrichment of neuronal growth factor (NGF) signaling, Insulin-like Growth Factor (IGF) regulation, cytokine signaling (IL-6, IL4 and IL13 signaling) and Role of PKR in Interferon Induction and Antiviral Response (Figure 3). These results show that repeated WS exposure induces time-dependent transcriptional changes that are associated with diverse biological processes in airway epithelial cells.

**Figure 3:**
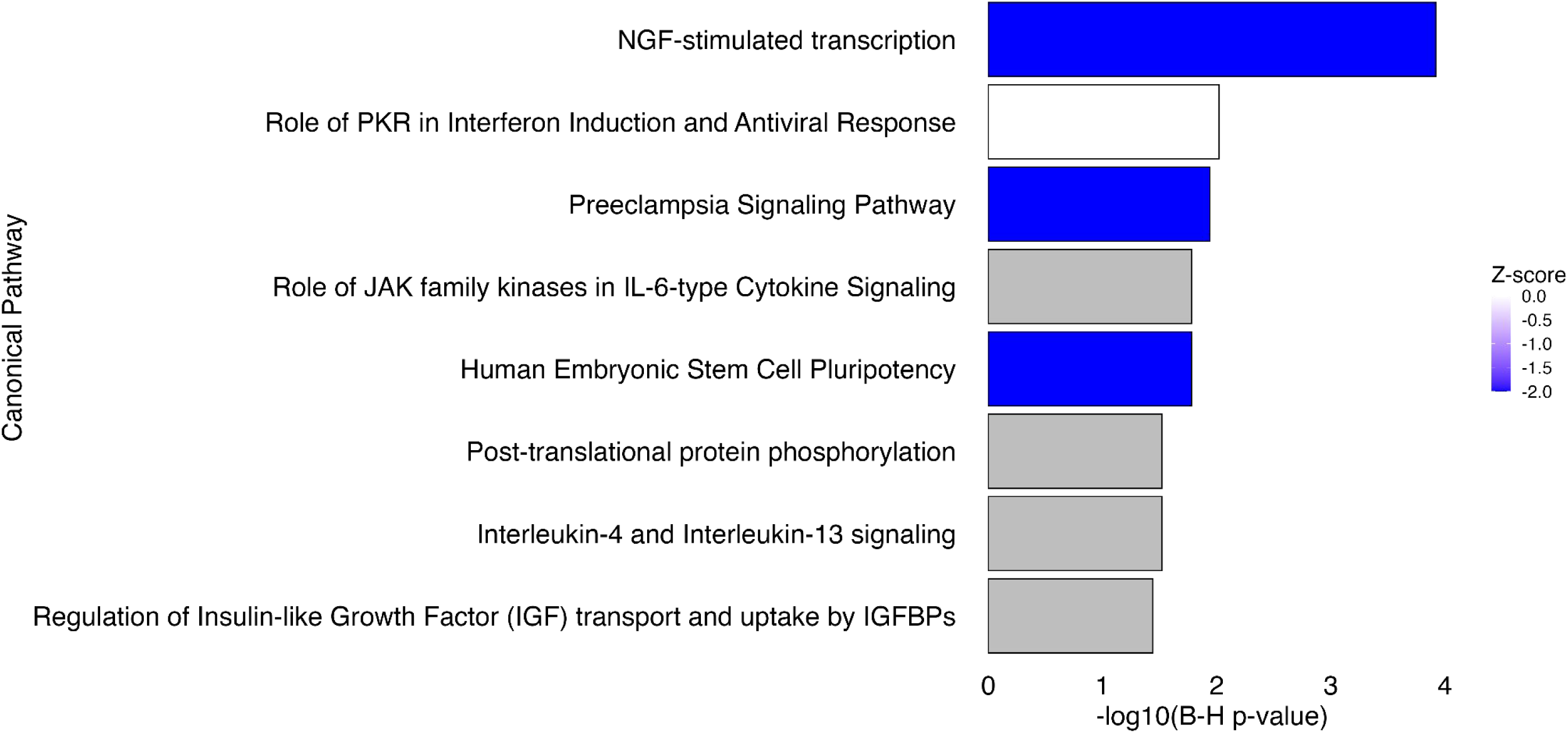
Pathways persistently affected by wood smoke exposure. Bar plot showing significantly enriched pathways from IPA of 77 DEGs shared between the 5-day and recovery timepoints. Z-scores are derived from differential expression of the 77 DEGs at the recovery timepoint. Overlapping DEGs between these two timepoints represent transcriptomic perturbations following repeated exposure to simulated WS that persist after one week of rest from exposure.

### Repeated Wood Smoke Exposure Induces Significant Changes in DNA Methylation

To assess the role of the epigenome in the observed dynamic transcriptional changes in airway epithelial cells following repeated WS exposure, we performed genome-wide DNA methylation analysis on adult rhesus airway epithelial cells following five days of exposure. 8,045 differentially methylated regions (DMRs) were detected (empirical p<0.05, difference in DNA methylation < = 5%) (Supplementary Table 3A), with 51% exhibiting hypermethylation in exposed cells (Figure 3A). Genomic distribution analysis revealed that these DMRs were significantly enriched in promoter and exon regions (p<0.05), while intronic regions were significantly depleted for DMRs (Figure 4B). Pathway analysis of genes most proximal to DMRs indicated significant enrichment for notable processes including xenobiotic metabolism, cytokine signaling, neuroinflammation signaling, pulmonary healing signaling, and regulation of the EMT by growth factors, highlighting potential regulatory mechanisms underlying the molecular response to WS exposure (Supplementary Table 4). Taken together, these changes in methylation underscores the impact of repeated WS exposure on pathways involved in toxicant detoxification, inflammation, and epithelial stress responses in airway epithelial cells.

**Figure 4:**
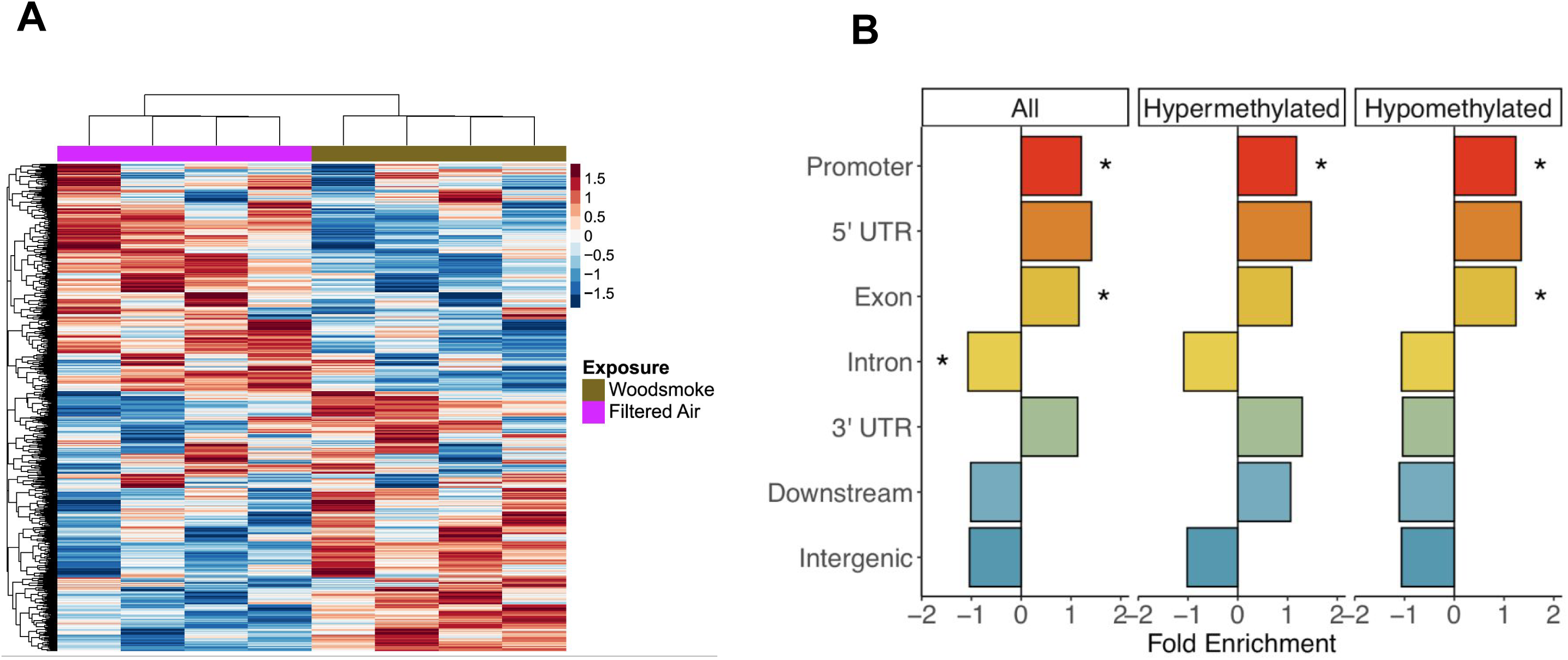
Repeated wood smoke alters DNA methylation in airway epithelial cells. A) Heatmap showing methylation of 8,045 DMRs across all samples following 5-day WS exposure. Red represents hypermethylation and blue represents hypomethylation. B) Bar graph showing enrichment and depletion of DMRs across genomic locations. The plot displays fold enrichment of DMRs across genomic features, separated into three groups: All, Hypermethylated, and Hypomethylated. The x-axis represents fold enrichment values, while the y-axis lists genomic features. Each colored bar corresponds to a feature’s enrichment value, and asterisks mark statistically significant categories.

### Correlation of DNA methylation and Gene Expression Differences Resulting From Repeated Wood Smoke Exposure

To determine the potential transcriptional consequences of these methylation changes, we integrated the genome-wide DNA methylation data with the transcriptomic profiles following repeated exposure. Fourteen percent of DEGs after five days of exposure were located close to DMRs (Supplementary Table 5A). Direct correlation analysis of percent difference in methylation and fold change values between overlapping DMRs and DEGs showed no significant linear relationship (r = -0.01, p = 0.928) (Figure 5A). Stratifying correlation analysis by DMR genomic locations strengthened correlation coefficients but did not reveal any significant relationships between methylation and expression changes after five days of exposure. As DNA methylation may cause persistent changes in gene expression, we also examined correlation between percent difference in methylation and fold change values between overlapping DMRs at 5-day exposure and DEGs detected after the recovery period and found no significant linear relationship (r = -0.057, p = 0.402, DEG and DMR pairs shown in Supplementary Table 5B).

**Figure 5:**
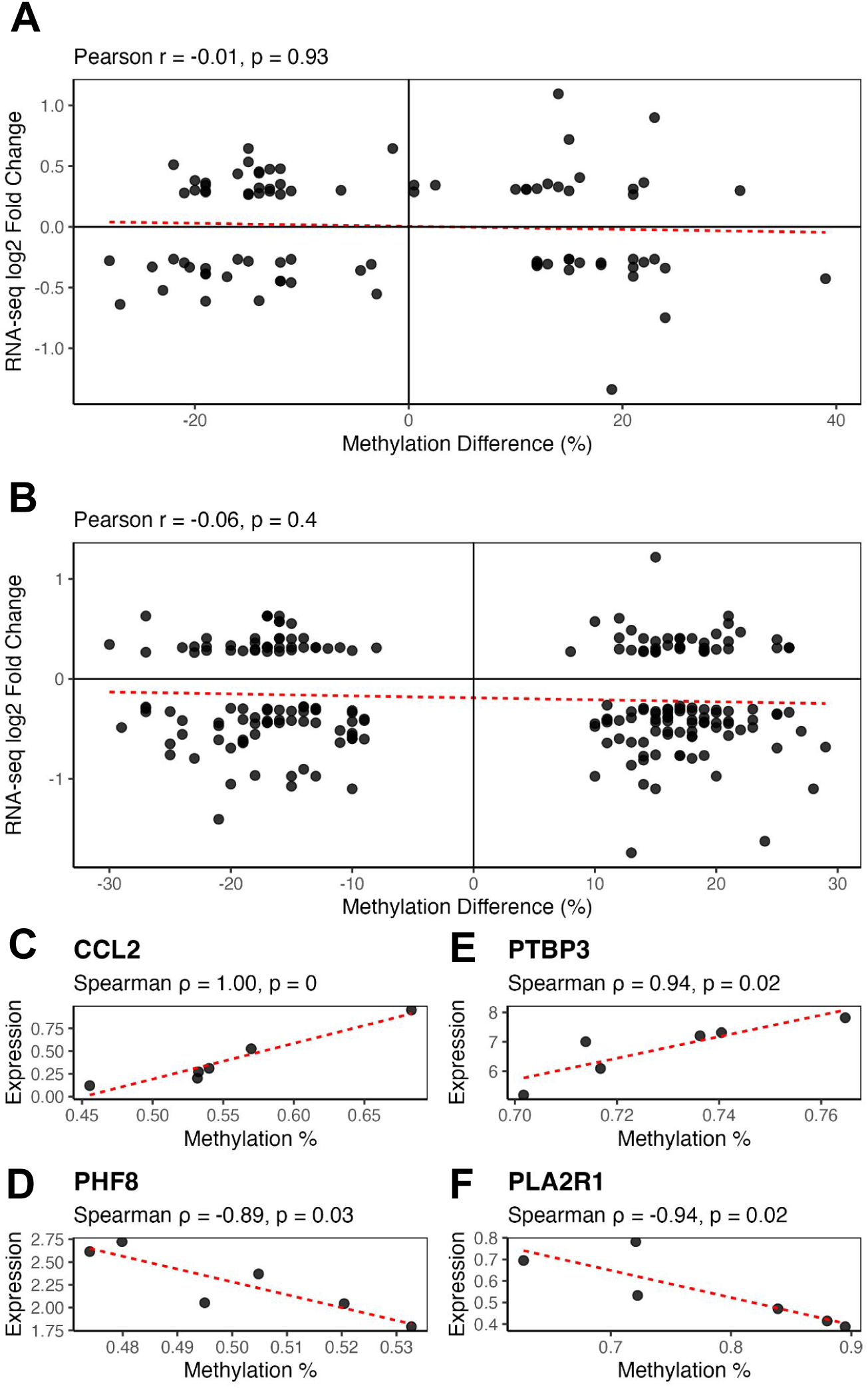
Correlations between DNA methylation and gene expression changes induced by wood smoke exposure. A) Global spearman correlations of percent methylation changes and gene expression fold changes following 5-day WS exposure. B) Global spearman correlations of percent methylation changes following 5-day WS exposure and gene expression fold changes following one week recovery from repeated WS exposure. C) Scatterplot showing spearman correlations of gene expression and DNA methylation levels across paired samples for CCL2. D) Scatterplot showing spearman correlations of gene expression and DNA methylation levels across paired samples for PHF8. E) Scatterplot showing spearman correlations of gene expression and DNA methylation levels across paired samples for PTBP3. F) Scatterplot showing spearman correlations of gene expression and DNA methylation levels across paired samples for PLA2R1. The line included in plots C-F represents the best-fit linear model line.

Among 8,045 DMRs, 2,904 had enough corresponding gene expression to evaluate the correlation between gene expression and DNA methylation. We identified 124 genes with significant correlations (p<0.05), even though most of these genes were not significantly differentially expressed (Supplementary Table 5C). Some of these genes were associated with lung disease and biological processes of interest such as cytokine signaling, chromatin regulation, and cell senescence (Figure 5C-F). Collectively, these results demonstrate that repeated WS exposure induces significant DNA methylation changes, with only a minority of these changes correlating with alterations in gene expression. This pattern suggests selective epigenetic transcriptional regulation and the involvement of distinct molecular mechanisms underlying the biological response to WS exposure.

### Wood Smoke Exposure Induces Persistent Effects on Gene Co-expression Networks

In addition to differential gene expression analysis, we applied weighted gene co-expression network analysis (WGCNA) to the RNA-seq data to investigate the broader transcriptional impact of WS exposure. This approach identified distinct modules of co-expressed genes associated with WS exposure across timepoints. The optimal soft threshold for the WGCNA was 17 based on the scale-free topological fit index (R2 = 0.8) (Figure 6A). A total of 16 gene modules were identified by hierarchical clustering (Figure 6B). The eigengenes of the darkorange, darkorange2, and lightyellow modules showed significant correlations with exposure conditions (Figure 6C). IPA of the genes in the darkorange module revealed significant enrichment for pathways involved in epithelial remodeling, inflammation/immune signaling, epigenetic regulation, and apoptosis/cell cycle control (Supplementary Table 6A and a subset are highlighted in Figure 7). The darkorange module showed significant negative correlation with WS exposure 5-day timepoint (Cor = -0.70, p = 2.2e−04) and the negative correlation remained but became non-significant (Cor = -0.27, p = 0.21) at the recovery timepoint (Figure 6D). IPA of the genes in the darkorange2 module returned pathways associated with epithelial remodeling and inflammation/immune signaling (Supplementary Table 6B). The darkorange2 module showed significant negative correlation with the WS exposure 5-day timepoint (Cor = -0.70, p = 2.2e−04) and a positive but not significant correlation arose (Cor = 0.22, p = 0.31) at the recovery timepoint (Figure 6D). IPA of the genes in the lightyellow module returned pathways associated with apoptosis/cell cycle control, proteostasis, immune/inflammatory activation, and mitochondrial metabolism (Supplementary Table 6C). The lightyellow module showed significant positive correlation with the WS exposure 5-day timepoint (Cor = 0.73, p = 9.1e−05). Among these three modules, genes with the most connections within the modules (hub genes) were identified and the top 25 were shown in Figures 6E-G. Taken together, the WGCNA analysis identified additional genes and pathways that are responsive and persistently impacted by WS exposure.

**Figure 6:**
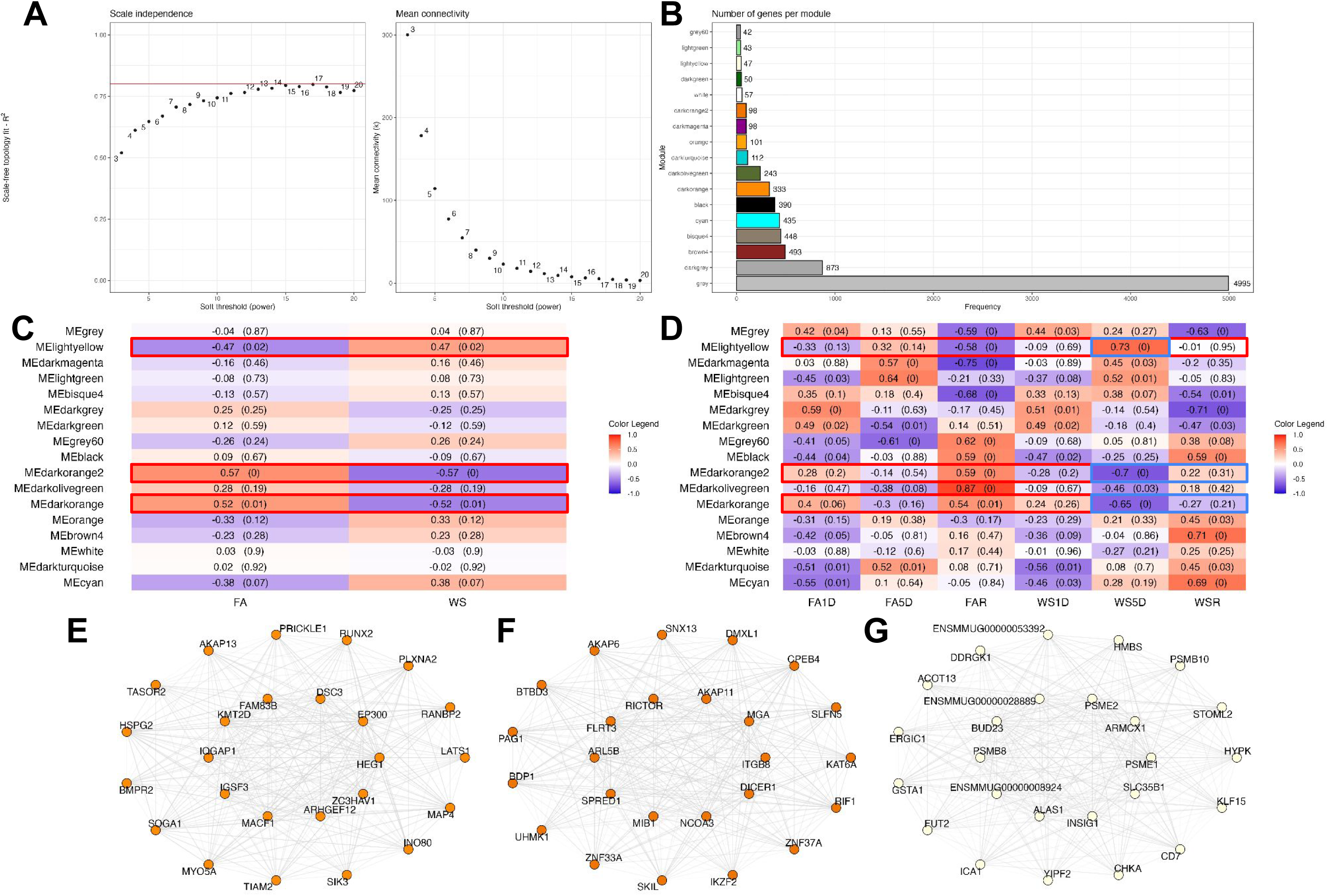
WGCNA analysis identified modules significantly correlated with wood smoke exposure. A) WGCNA soft-thresholding analysis showing power selection for network construction (β = 17). B) Bar graph showing the number of genes in each gene module from WGCNA. Sixteen gene modules were identified. C) Heatmap of module-exposure relationships showing eigengene correlations with exposure timepoints. In each cell, Pearson correlations between eigengene expression and exposure types are featured with corresponding significance values in parentheses. D) Heatmap of module-exposure relationships stratified by exposure timepoint. Modules with significant correlations with WS exposure are highlighted with red boxes in C and D. E) Connectivity plot of the top 25 darkorange module hub genes. F) Connectivity plot of the top 25 darkorange2 module hub genes. G) Connectivity plot of the top 25 lightyellow module hub genes. Hub genes were ranked and selected for plotting based on gene expression Pearson correlation with the ME. The top 10 ranked hub genes were plotted within the remaining 15 hub genes.

**Figure 7:**
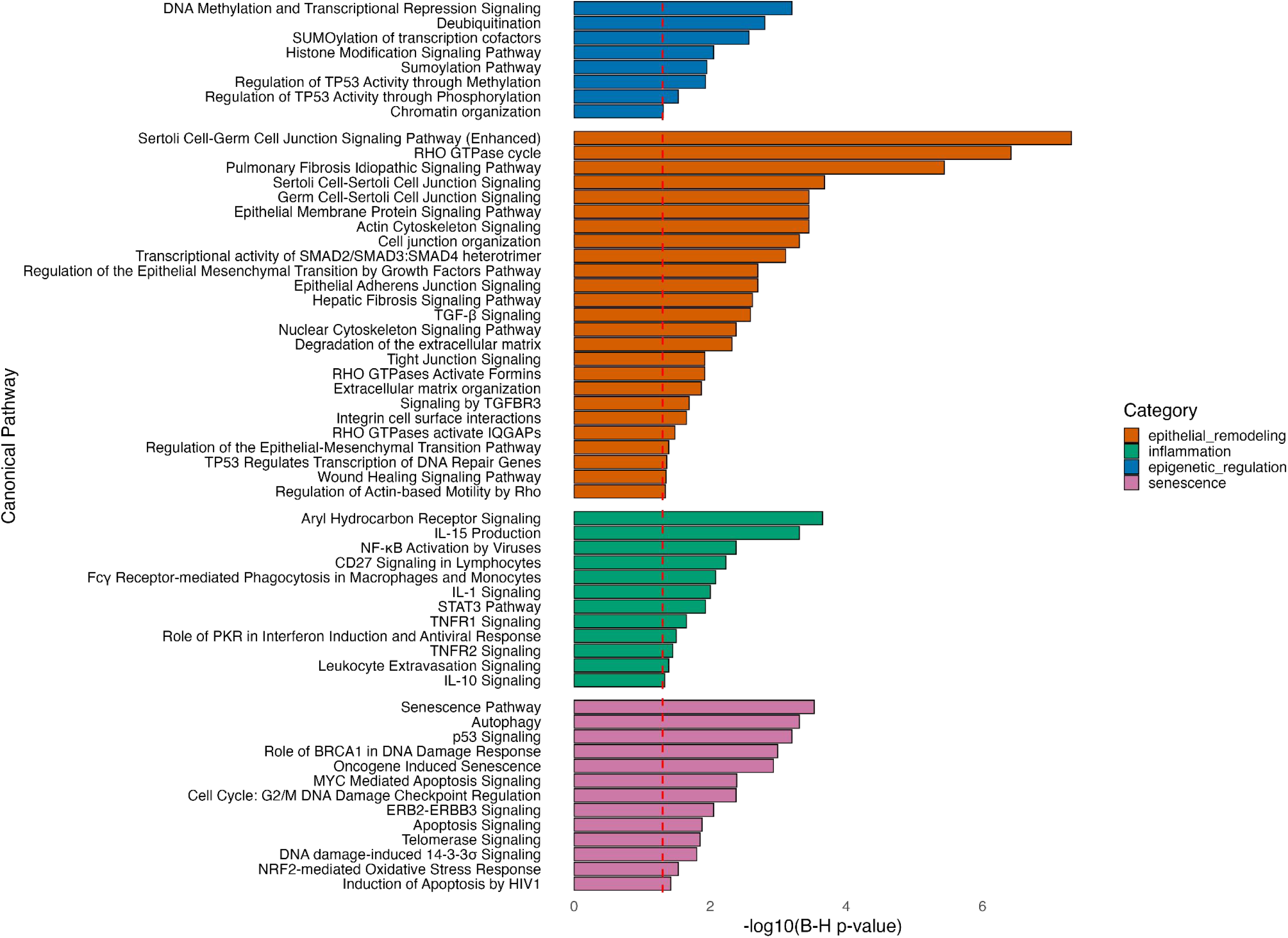
Bar graph of IPA results for the darkorange module genes. Pathways of interest were selected from significantly enriched pathways for plotting by match with words or phrases in a manually generated key words list for terms relevant to senescence, epithelial remodeling, inflammation, and epigenetic regulation.

### Wood Smoke Induced Gene Expression Perturbations Overlapped with Lung Disease Signatures

Finally, we assessed the similarity between transcriptomic exposure signatures and publicly available human lung and nasal epithelial transcriptomic signatures for respiratory diseases using a previously established method (Koval et al., 2022). DEGs following 1 day of WS exposure had low similarity to both five days of exposure (Jaccard index=0.02, 11 shared genes) and recovery DEGs (Jaccard index=0.01, 5 shared genes), while DEGs following five days of exposure and recovery shared the highest Jaccard index among the exposure groups (0.06, 74 shared genes) (Figure 8A). Between disease signatures, COPD and IPF DEGs exhibited the highest inter-disease similarity (Jaccard index=0.25). Asthma showed minimal DEG overlap with the other diseases (Jaccard index=0.04). WS exposure showed limited overlap with represented asthma, COPD, and IPF signatures (≤ 0.02), but DEGs following recovery had a slightly higher overlap with COPD (Jaccard index= 0.02, 292 shared genes) and IPF (Jaccard index = 0.02, 354 shared genes) than the other exposure groups. As these small indexes are partially due to the large number of DEGs associated with diseases, especially for COPD and IPF, we further evaluated the proportion of disease signatures in the exposure signatures and found that about half of the DEGs following five days of exposure and recovery were represented by COPD and/or IPF DEGs (Figure 8B). The exposure signature genes represented in both the COPD and IPF signatures were significantly enriched for pathways involved in inflammation, growth factor signaling, epigenetic regulation (Figure 8C). Collectively, these comparisons linked WS induced, time-dependent transcriptomic changes with many pathways featured in lung diseases.

**Figure 8:**
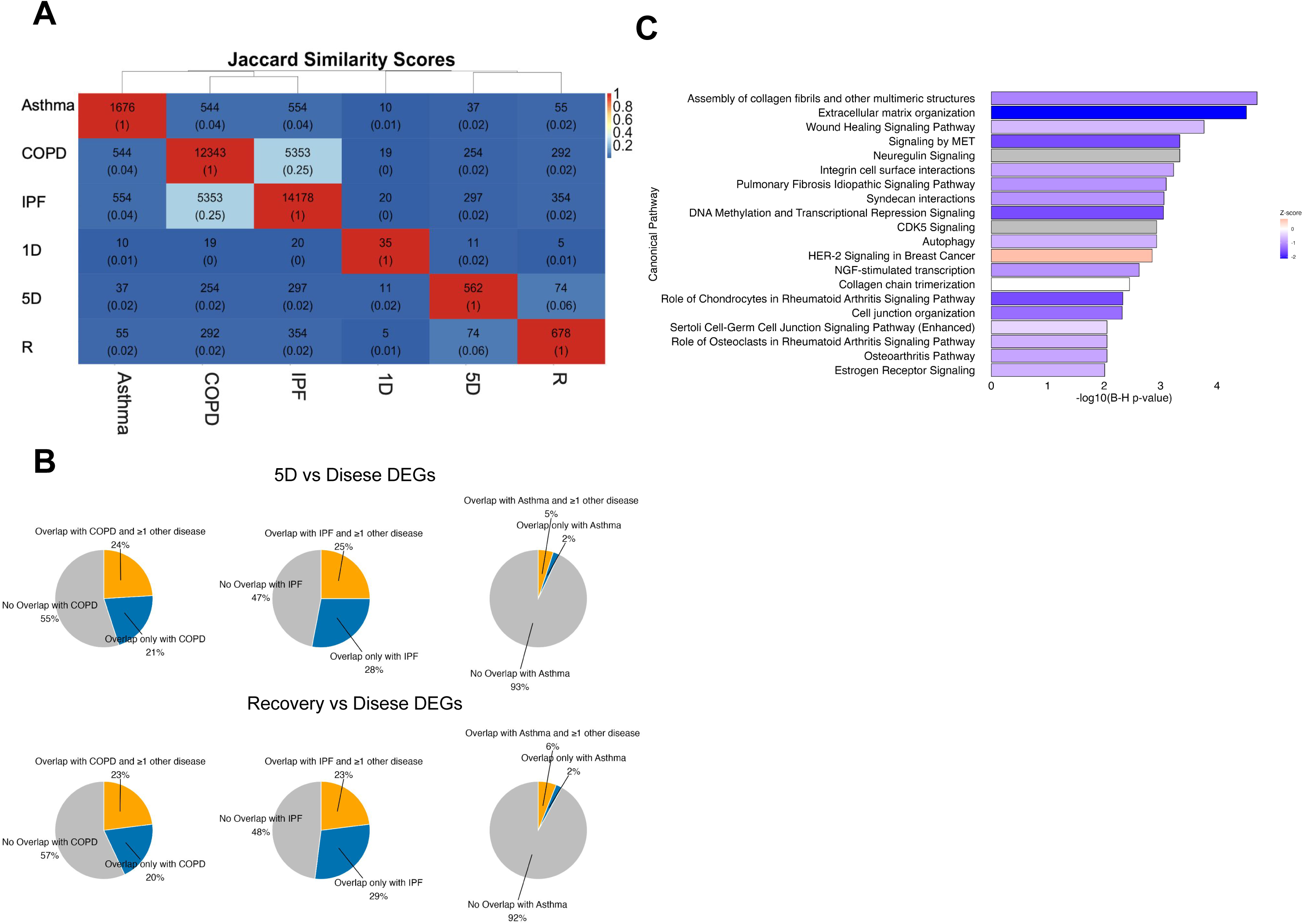
Enrichment of respiratory disease-associated gene expression signature in wood smoke exposure induced transcriptomic changes. A) Heatmap of Jaccard similarity indices for DEGs across exposure and disease groups. In each cell, top values represent the intersect between DEGs of the corresponding transcriptomic profiles. Bottom values represent the ratio of the intersect of DEGs over the union of the DEGs of the corresponding transcriptomic profiles. B) Pie charts of the proportion of DEGs from exposure groups that overlap with COPD, IPF, and asthma disease signatures. Grey wedges represent the proportion of exposure DEGs that are unique to the corresponding exposure condition. Blue wedges represent the proportion of DEGs that overlap between the corresponding exposure condition and the corresponding disease signature. Orange wedges represent the proportion of DEGs that overlap between the corresponding exposure condition and multiple disease signatures. C) Bar graph showing broad overlap in enriched biological processes between persistent transcriptomic changes following repeated WS exposure and COPD and IPF. IPA results for genes overlapping all four sets are plotted: COPD signature, IPF signature, 5-day exposure, and recovery DEGs.

## Discussion

### Dynamic and Persistent Transcriptomic Changes After Wood Smoke Exposure

Our findings demonstrated that repeated WS exposure in differentiated adult rhesus macaque airway epithelial cells provokes robust changes in gene expression, with both dynamic and persistent elements. Most strikingly, a surge of transcriptomic alterations arose after 5 days of repeated exposure. Ferroptosis signaling was enriched as a significantly upregulated pathway. Parallel exposures (PM2.5 and cigarette smoke) in BEAS-2B and primary epithelial cells showed lipid ROS accumulation, GPX4 depletion, SLC7A11 inhibition, and ACSL4-linked phospholipid remodeling rescued by ferroptosis inhibitors or NRF2 activation (Ahn et al., 2024, Li et al., 2025, Qin et al., 2024, Zhao et al., 2025). These data suggest a significant interaction between chemical features of WS (oxidants, quinones, metals) and ferroptosis signaling that may contribute to epithelial lipid peroxidation, barrier failure, and sustained inflammatory signaling.

The number of differentially expressed genes increased further following a 7-day recovery period (Supplementary Table 1). Unlike at other timepoints, expression changes indicated the downregulation of pathways involved in tissue repair and remodeling (such as those regulating growth factor signaling, extracellular matrix production, or cell adhesion). The darkorange module was the largest gene module significantly associated with WS exposure, and featured enrichment in pathways related to inflammation, wound healing, and tissue remodeling (Figure 6B-C & Figure 7). This module showed a sustained expression pattern where genes were downregulated after 5 days of repeated exposure and remained suppressed after a one-week recovery period only in the WS exposed samples (Figure 6D). These sustained transcriptional alterations point to a longer-lasting impact of WS on the airway epithelium that extends beyond the exposure period. Such an impact of repeated smoke exposure on the airway epithelium could cumulatively lead to lasting changes in epithelial biology, potentially priming the airway epithelium for the development of chronic respiratory issues.

In parallel with gene expression profiling, our study revealed concurrent changes in the epigenome of airway epithelial cells following WS exposure. We identified several differentially methylated regions (DMRs) following repeated exposure, reflecting epigenetic remodeling in response to WS. Notably, only a minority of genes showing expression changes also featured significant shifts in nearby methylation, and these transcriptomic and epigenetic changes generally presented no significant correlations (Supplementary Table 5C). This apparent disconnect suggests that transcriptional responses and DNA methylation changes may operate on different timescales or involve niche sets of genes. In this context, DNA methylation alterations might represent a longer-term mechanism of regulation that could stabilize gene expression changes or prime certain loci for future responses. The disconnect we observed underscores the intricate multi-level response of airway cells to inhaled smoke that warrants further investigation.

Some genes nearest to DMRs, with significantly correlated expression and methylation, were involved in chromatin remodeling and airway inflammation associated processes. Our results suggest that the genes involved in these processes influence one another in driving tissue responses and disease outcomes. Among these genes, C-C motif chemokine ligand 2 (CCL2) is a chemoattractant for monocytes that may act as a central mediator of the inflammatory response to WS exposure (Figure 5C). Multiple studies indicate that CCL2 is upregulated upon biomass smoke extract exposure. *Ex vivo* experiments with human lung tissue exposed to biomass particulate matter showed significant increases in CCL2, as well as with other inflammatory cytokines like IL-1β, IL-6, IL-8, TNF-α and chemokines CCL3 and CCL13 (Kc et al., 2020). Elevated CCL2 in the airway epithelium may contribute to recruiting monocytes/macrophages to the lungs, which produces more inflammatory mediators and potentially exacerbates tissue injury as observed in mouse models (Pardo et al., 2020). Furthermore, a recent review underscored that inflammation, oxidative stress, and cytotoxicity are key impacts of wildfire smoke on airway epithelial cells that lead to lung damage (Heibati et al., 2025).

Notably, several chromatin-modifying genes associated with DMRs had significantly correlated expression and methylation. One such gene is PHF8, a Jumonji family histone demethylase that typically removes repressive histone methylation marks (such as H3K9me2, H3K27me2, and H4K20me1) to activate transcription (Qi et al., 2010) (Figure 5D). In human lung cells, PHF8 was identified as a factor associated with smoking-related lung cancers (El-aarag et al., 2017). Many histone-modifying enzymes like PHF8 are sensitive to cellular redox and metabolic state. PHF8’s demethylase activity requires molecular oxygen and iron as cofactors, so oxidative stress from WS could affect its function (Fan et al., 2024). By altering methylation marks on chromatin, PHF8 could change transcription of genes involved in stress responses, cell cycle, or inflammation. If wood smoke exposure changes the expression of chromatin modifiers, it could epigenetically perturb certain inflammatory genes, so that subsequent exposures may elicit an altered response, even if the expression of these genes is not immediately changed.

### Integration with Previous Early-Life Exposure Findings

This study extends our prior work examining the long-term epigenetic effects of early-life wildfire smoke exposure in rhesus macaques (Brown et al., 2021). In our previous early-life exposure model, animals exposed to ambient wildfire smoke as infants showed thousands of persistent DNA methylation changes in their nasal epithelium years later. Strikingly, minimal lasting gene expression differences were observed. This suggested that early-life smoke exposure triggers extensive epigenetic reprogramming without overtly altering baseline transcriptomes in the long term. Those latent epigenetic alterations were enriched at genes involved in immune and neural system development, including regions of bivalent chromatin that regulate developmental gene expression. In contrast, airway epithelial cells from the adult animals in our study exhibited pronounced transcriptional responses to WS, potentially reflecting the difference in response between exposure times (Supplementary Table 1). The gene expression changes we observed (inflammation, stress, and repair genes) in this study were largely unobserved in our early-life cohort at baseline, likely because those animals were examined long after the exposure.

However, we found that both early-life and repeated adult exposure led to significant DNA methylation changes. There were 726 shared genes associated with DMRs in both studies; these genes are involved in processes such as development, oxidative stress response, and immune response. Additionally, there were 50 positional overlaps amongst the DMRs from the two studies (Supplementary Table 3B). Altogether, these transcriptomic and epigenomic observations suggested that the effects of WS on gene expression persist days after exposure, but most changes may resolve into epigenetic memory with prolonged recovery time. This also demonstrates that epigenetic plasticity in the airway extends beyond early developmental windows into adulthood. However, the context and consequences of these epigenetic changes may differ from early-life exposures. Whereas early-life smoke may disrupt developmental epigenetic programming (potentially affecting how the immune system matures or how the airway structure develops), adult exposure-induced methylation changes might influence the function of mature cells or the activity of tissue-resident progenitor cells involved in repair. This means that detecting DNA methylation changes in cells could be indicative of past wildfire smoke exposure and a propensity for altered responses, which is supported by a recent study of blood DNA methylation profile associated with chronic exposure to wildfire-derived particulate matter in humans (R. Xu et al., 2023). In the context of our findings, it is possible that the methylation changes observed in the airway epithelium after WS exposure could become long-term memory and prime the tissue for an exaggerated or dampened response to subsequent insults, which warrants future investigation. Taken together, these findings support a model in which epigenetic changes in response to WS may both accompany active gene regulation and serve as a persistent memory of exposure that might modify future gene expression programs.

### Co-expression Network Analysis Reveals Coordinated Epithelial Response

To better understand the coordinated biological processes activated by WS exposure, we employed WGCNA. This approach clustered genes into modules based on similar expression patterns across the time course, offering insight into networks of genes that act in concert. Genes in the previously mentioned darkorange module showed a persisting downregulation response induced by WS (Figure 6C). Among these genes, we found enrichment for inflammatory signaling pathways, such as those involved in IL-10 signaling and leukocyte extravasation (consistent with recruitment of immune cells to the airway) and other cytokine and chemokine networks (Figure 7). These genes were also enriched for pathways related to oxidative stress and xenobiotic metabolism, including AhR signaling (Figure 7). Additionally, the AhR gene was featured among the darkorange2 module genes (Supplementary Table 6B). AhR is a key sensor of environmental toxins like polyaromatic hydrocarbons present in smoke, and activation of AhR-regulated genes (e.g., detoxification enzymes and phase I/II metabolism genes) is a well-known response to inhaled pollutants (Ho et al., 2022). The presence of AhR pathway genes in these modules aligns with known responses to particulate matter and organic components of WS that can engage AhR and underscores possible perturbations in xenobiotic metabolism with negative impacts on oxidative stress and inflammation.

In addition to inflammatory and xenobiotic response pathways, the darkorange module genes, whose downregulated expression persisted following repeated exposure and the recovery period, were enriched in pathways indicative of epithelial injury and repair. For example, we observed over-representation of signaling cascades involved in cell adhesion, junction remodeling, and cytoskeletal dynamics (e.g., Rho GTPase effector pathways and integrin signaling) (Figure 7). Pathways related to tissue remodeling and fibrosis, such as the idiopathic pulmonary fibrosis signaling pathway and EMT regulation by growth factors, were significantly enriched (Figure 7). The darkorange module gene encoding Polypyrimidine Tract Binding Protein 3 (PTBP3) has been observed to drive EMT and invasion in lung adenocarcinoma via the TGF-β/Smad pathway (Dong et al., 2022). PTBP3 was not detected in this study to be significantly differentially expressed following WS exposure, but it featured a hypermethylated intronic region and had significantly positively correlated expression and methylation across samples (Supplementary Table 3A and Figure 5E). Impairments to wound healing or chronic injury response may be introduced with the downregulation of the genes involved in these pathways, as observed in the darkorange module. Furthermore, the darkorange module was enriched for senescence pathways related to tissue remodeling and aging (Figure 7). The darkorange module gene encoding M-type Phospholipase A₂ Receptor 1 (PLA2R1) has been observed to mediate a senescence-associated secretory phenotype (SASP) in COPD (Beaulieu et al., 2021). While chronic SASP is associated with tissue pathology, transient SASP has been observed to contribute to wound healing (Wilkinson & Hardman, 2020). PLA2R1 was detected in this study to have reduced expression in WS exposed samples following the one-week recovery period (Supplementary Table 1C), featured a hypermethylated intronic region and significantly positively correlated expression and methylation across all samples (Supplementary Table 3A and Figure 5F). Suppression of SASP, characterized by decreased pro-inflammatory and pro-fibrotic signaling, in airway cells may contribute to insufficient initiation of repair mechanisms after injury. This shift toward reduced wound healing processes was reflected in the downregulation of key regulatory genes such as SMAD3, TP53, and REL.

Comparison with other modules underscores the dynamic interplay between injury, inflammation, and remodeling. The darkorange2 module was similarly suppressed under repeated WS exposure and was significantly enriched for IL-6 family signaling and JAK/STAT cascades, along with IL-27 signaling (Supplementary Table 6B). IL-6 family and IL-27 signaling in airway epithelial cells are known to activate JAK/STAT pathways for regulating immune responses, epithelial homeostasis, and dampening chronic inflammation and EMT (Abaurrea et al., 2021). Suppression of these potentially protective signaling pathways may result in reduced anti-inflammatory activity, impaired epithelial recovery, or unchecked remodeling/fibrosis when exposed to chronic environmental stress. By contrast, the lightyellow module was strongly upregulated with repeated exposure and enriched for oxidative stress, non-canonical NF-κB signaling, and apoptosis-related processes (Supplementary Table 6C). This reflects an injury and inflammatory program that dominated while remodeling pathways were repressed. Together, these observations indicate that WS drives a temporal shift in airway epithelial state where structural remodeling programs represented by the darkorange and darkorange2 modules are suppressed while injury/immune programs represented by the lightyellow module are amplified following repeated WS exposure.

Growth factor signaling pathways associated with darkorange and darkorange2 module genes may be central to immune-repair pathway interactions. Interestingly, the top enriched IPA pathway from genes among the overlapping genes with perturbed expression following 5 days of repeated exposure and a 7-day recovery period was NGF signaling (Figure 3). Pathway enrichment analysis of the DMRs also detected perturbations in neuroinflammation signaling (Supplementary Table 4). Furthermore, darkorange WGCNA module genes were enriched for NGF signaling and several signaling cascades downstream of NGF stimulation (Supplementary Table 6A). Canonical pathways involved in NGF signaling via MAPK, ERK5, and RhoA were also detected to be reduced persistently into the one-week recovery period following repeated WS exposure. These pathways are known mediators of stress-activated cellular responses, including inflammation, cytoskeletal remodeling, and survival/apoptotic balance (Xing et al., 1998; Nusser et al., 2002; Wang et al., 2006). Our observation that repeated WS exposure attenuates expression of genes within these NGF-regulated pathways suggests that WS may interfere with the capacity of airway epithelia to regulate repair and survival programs. This is consistent with existing evidence that NGF stimulation in bronchial epithelial cells promotes anti-apoptotic activity and supports cellular proliferation (Othumpangat et al., 2009).

In summary, the co-expression of genes in the darkorange, darkorange2, and lightyellow modules suggests that pathways of inflammation (NF-κB/REL), cell stress and senescence (p53), and tissue repair signaling (TGF-β/SMAD, NGF/NGFR) are simultaneously perturbed in the airway epithelium. This indicates that the transcriptomic response to repeated WS exposure is characterized by networks that drive inflammation, oxidative stress, and cell survival while compromising barrier repair and differentiation, changes that may contribute to chronic airway remodeling and pathology.

### Implications for Chronic Respiratory Disease and Translational Relevance

The sustained molecular changes observed in our study have important implications for long-term respiratory health. Even though our experimental timeline was limited to one-week post-exposure, the persistence of perturbed inflammatory and remodeling gene expression programs at day 7 of recovery suggests that the cells did not fully return to a homeostatic state in that time. If such smoke exposures are recurrent, as in the case for wild animals or humans living in wildfire-prone regions, or firefighters experiencing repeated smoke inhalation, the incomplete recovery between exposures could lead to cumulative damage. Chronic disruptions of pathways like TGF-β/SMAD (wound healing), NF-κB (inflammation), and senescence can drive pathological remodeling of the airway wall, susceptibility to pathogens and harmful pollutants, and loss of normal epithelial function (Zaynagetdinov et al., 2015). In support of this, one week after repeated WS exposure, the transcriptomic changes we observed mirrored features observed in chronic respiratory diseases such as COPD or IPF (Figure 8). Our finding that transcriptomic changes were enriched in an idiopathic pulmonary fibrosis signature was noteworthy. It raised the possibility that the observed effect of repeated WS inhalation on gene expression might invert to push the airway toward a fibrosis-like phenotype, especially in susceptible individuals or alongside other risk factors. Furthermore, impairments in epithelial repair and regeneration, may reduce the airway’s ability to resolve injury, leading to persistent epithelial disruption and inadequate wound healing. These mechanisms are central to the development and progression of COPD. While histology staining did not suggest morphological changes (Supplementary Figure 1), the gene expression patterns provided molecular evidence to support epidemiological links between WS exposure and respiratory morbidity, including increased hospital visits for asthma exacerbations, COPD flare-ups, and other respiratory complications (Zhang et al., 2025). Our data offered a mechanistic glimpse into how such exacerbations might arise at the molecular and cellular level.

Our results also highlight the translational relevance of using rhesus macaques as a model for inhaled smoke exposure. Nonhuman primates (NHP) share many anatomical, cellular, and immunological features of the human respiratory system, making them especially informative for understanding complex exposures like wildfire smoke and establishing causal link between exposures and pathological changes (Miller et al., 2017). The agreement of pathways identified in our *in vitro* studies using rhesus airway cells with those reported in human studies (inflammation, oxidative stress, etc.) suggest that we observed fundamental biological responses that likely occur in exposed humans as well (Zhai et al., 2022). This adds confidence that interventions targeting these pathways in future NHP *in vivo* models could be beneficial in humans. For instance, the prominence of oxidative stress pathways (e.g., AhR and ferroptosis signaling) in the airway response might validate ferroptosis inhibitor or AhR antagonist strategies to mitigate smoke-induced damage. A clear implication of our findings is that time for recovery is crucial. Specifically, even after cessation of exposure, the airway epithelium may need more than a week to fully recover. Repeated wildfires or prolonged smoke seasons could shorten these recovery windows, compounding the effects. Therefore, public health strategies during wildfire events might consider not just acute exposure reduction but also post-exposure monitoring and therapeutic support to ensure proper recovery of the respiratory system.

## Limitations and Future Directions

While our study provided valuable insights into the transcriptional and epigenetic effects of repeated WS exposure, several limitations warrant consideration. First, our observations are from *in vitro* differentiated airway epithelial cells from adult female animals. Sex-specific susceptibility to environmental exposures has been reported in the development and progression of respiratory diseases such as asthma and COPD, particularly for women (Fuentes et al., 2017, Calzetta et al., 2025, Ober et al., 2008). Future studies will determine whether similar responses to WS in airway epithelial cells occur *in vivo*, and how age and sex influence these responses. Second, our study focused exclusively on airway epithelial cells, which do not capture the contributions of other critical cell types such as immune cells, fibroblasts, or endothelial cells. Future single-cell or spatial transcriptomic approaches could clarify the cell-type-specific nature of these responses and reveal interactions between epithelial, vascular, structural, and immune compartments during and after exposure. Third, we used simulated WS to expose the cells, which is biomass burning and similar to wildfires primarily occurring in the wildland. Other types of wildfires, such as those that occur at wildland-urban interface, contain structure burning and have additional toxic components such as metals that warrants further investigations (Baliaka, H. D., Ward, R. X., Bahreini, R., et al., 2025 and California Air Resources Board, 2025).

Two additional limitations were the short follow-up period for transcriptomic analysis and single time point for methylation analysis. Although we sampled animals for transcriptomic analysis up to 7 days post-exposure, this window may not fully capture the resolution or persistence of molecular changes. This limitation was magnified for the observation of epigenetic modifications, which can evolve over weeks or months. Longer-term studies are needed to determine whether the observed methylation changes persist, whether gene expression returns to baseline, and how these molecular patterns relate to functional outcomes such as barrier integrity, tissue remodeling, or susceptibility to secondary insults (e.g., viral infections).

Furthermore, while our integrated analysis examined correlations between DNA methylation and gene expression, causality remains unresolved and mechanisms other than DNA methylation may be involved. Functional validation, such as CRISPR-mediated epigenome editing or reporter assays, will be essential to determine whether specific methylation changes are sufficient to drive transcriptional regulation and provide critical insights into the design of precision epigenetic therapeutics.

## Conclusion

We demonstrated that repeated wood smoke exposure elicits dynamic transcriptional responses, coordinated gene network remodeling, and widespread epigenetic alterations. Persistent gene expression changes following a short recovery period suggested incomplete recovery and disruption of pathways related to wound healing, senescence, and epithelial remodeling, which are processes involved in chronic respiratory disease development. Despite the identification of thousands of differentially methylated regions, gene expression and DNA methylation changes were largely uncoupled, indicating distinct regulatory mechanisms operating across different timescales. These findings extended our previous work on early-life exposures and highlighted the complexity of airway epithelial responses to repeated smoke inhalation. Together, these data underscored the potential for WS to leave a lasting molecular imprint on the airway epithelium, with important implications for respiratory health in populations exposed to wildfire events.

## Methods

### Rhesus macaque primary tracheobronchial epithelial cell culture

Tracheobronchial specimens from 5 female rhesus macaques (3, 6, 6, 8, 14 yrs of age) were obtained from the California National Primate Research Center Pathology Service and airway epithelial cells were isolated as described by Wu, et. al. (Wu et al. 1986) and cryopreserved until use. Primary airway epithelial cells were expanded in two passages before plating to 12mm Transwell (0.4m pore size polyester, Corning, Inc.) inserts in a 12 well format, previously coated with FNC Coating Mix (Athena Enzyme Systems). Once visually confluent, apical media was removed for subsequent culture at an air-liquid interface (day 0 of ALI culture). Cultures from one macaque (6yr old) would not progress to ALI culture, limiting the experiments to four individuals. During ALI culture an apical wash with warm HBSS was conducted every 3-4 days until the exposure period. Initial expansion was supported by BEGM media (Promocell GmbH) and ALI culture in BEGM:DMEM (50% each v/v) supplemented with 50nM retinoic acid with media changes every 2-3 days.

### Biomass combustion product generation and exposures

The simulated WS consisted of biomass in the form of commercially available Douglas fir wood pieces cut to 1.9cm square by 61cm in length and combusted at 750° C. The biomass was fed into a quartz tube furnace at a constant rate of approximately 3g/min along with 10 L/min filtered air. Resulting combustion products were fed into a 4.3m^3^ chamber with dilution by HEPA and activated charcoal filtered air at a rate of 15 chamber volume changes per hour. The combustion products were metered by partial diversion directly to exhaust based on proportional feedback control to maintain a target particulate matter (PM) concentration of 75 μg/m^3^ the first day and 150μg/m^3^ the following four days. Chamber PM concentration was continuously monitored with a light scatter laser photometer (DustTrak model 8520) calibrated to the combustion PM by gravimetric analysis of PM samples from the combustion system, which served as the input for the proportional feedback control system. Carbon monoxide was monitored in the chamber by a gas filter correlation carbon monoxide analyzer (Teledyne API, model 300E). Chamber aerosol was sampled throughout each exposure day on filters (Pall Emfab TX40HI20WW) and analyzed gravimetrically for mass concentration. On the final day of the five-day exposure period, particle sizing was done at 1, 3 and 5 hours of the 6-hour exposure with a laser optical particle counter (Climet CI500). Chamber particle counting was conducted in bins of 0.3-0.5, 0.5-1, 1-5, 5-10, 10-25, and >25μm particle size. The chamber air mixture was drawn at 17 L/min through a temperature-controlled humidifier to achieve >95% relative humidity and carbon dioxide was added at 5% before entry into a 26.7L *in vitro* exposure chamber within an incubator maintained at 37° C.

Macaque primary airway epithelial cells cultured at air-liquid interface for 35 days were exposed to either filtered air or the combustion product mixture for either 1 or 5 consecutive days of 6hr exposure/day. Exposure of the single exposure group was conducted along with day 2 of the 5-day exposure group when the target PM level was 150μg/m^3^. A subset of cells undergoing 5 days of exposure also underwent one week of recovery in a standard cell culture incubator at 37° C, 100% relative humidity, and 5% carbon dioxide.

### Airway Epithelial Histology

A subset of cultures from each exposure group were fixed in Carnoy’s solution immediately after the last exposure. Following overnight fixation, cells were stored in 100% ethanol until paraffin embedding and sectioning as in Manna, et.al. (Manna, et. Al. 2021), which provides pairs of sections across the middle of the transwells. Sections were sent to UC Davis Veterinary Medicine Teaching Hospital (VMTH) histology laboratory for combined alcian blue and periodic acid-Schiff stains.

### Sample Collection & DNA/RNA Extraction

Rhesus macaque primary tracheobronchial epithelial cells were lysed using buffer RLT from AllPrep DNA/RNA micro kit (Qiagen, Hilden, Germany) supplemented with beta-mercaptoethanol (Bio-Rad, Hercules, CA) and was incubated at room temperature for 30 minutes. The cells were carefully scraped off the transwell membrane using a pipette and was aspirated to a microtube and was vortexed for 10 seconds. A Qia-shredder spin column (Qiagen, Hilden, Germany) was used to homogenize the cells where they were centrifuged for 2 minutes at full speed. The lysate from the column was transferred to a DNA spin column and was centrifuged for 30 seconds at >8000 x g. The DNA column was saved and placed on ice while the lysate was used to extract RNA after 70% ethanol was added and mixed. RNA was extracted using the RNA spin column following the manufacturer’s protocol. DNA was extracted from the DNA spin column following the manufacturer’s protocol. Qubit 4 fluorometer (ThermoFisher Scientific, Waltham, Massachusetts) was used to quantify both RNA and DNA.

### Library preparation for RNA sequencing (RNA-seq)

RNA (0.5µg per sample, all with RNA quality ≥ 7) was submitted for poly-A RNA library preparation and sequencing at Novogene on a NovaSeq 6000 (Sacramento, CA). Each sample had 20-33 million paired-end, 150 base pair reads (Supplementary Table 7A). RNA-seq read quality was assessed using FastQC. Trim Galore was used to trim bases with an adapter stringency set to 6, bases with less than a quality score of 20, 10 bases from the 5’ end of both read 1 and read 2, and reads smaller than 50 bases after trimming were discarded. Trimmed reads were aligned to the Rhesus monkey transcriptome (Mmul10) using Bowtie2 and quantified using RSEM (Langmead & Salzberg, 2012 and Li & Dewey, 2011). Transcript counts were subsequently converted into DESeq2 format using tximport for downstream analysis (Soneson et al., 2015).

### Differential gene expression analysis

Differential expression analyses were performed by DESeq2 (Love et al., 2014). Genes with an absolute shrunken fold change of at least 1.2 and an FDR ≤ 0.05 were considered significantly differentially expressed. Non-parametric correlation analysis using the Spearman rank correlation coefficient was used to examine the relationship between differential methylation and gene expression changes as well as normalized expression counts and methylation percentages. Specifically, percent methylation and percent methylation differences at genomic regions were averaged by shared gene annotations. These averaged values were correlated with corresponding gene or DEG expression counts and log₂ fold changes, respectively. Pathway analyses were performed using IPA (QIAGEN Inc., https://www.qiagenbioinformatics.com/products/ingenuitypathway-analysis) with an FDR ≤ 0.05 significance threshold.

### Weighted gene co-expression network analysis

A treatment group co-expression network was constructed using the WGCNA R package (Langfelder & Horvath, 2008). Genes with variance in the 25th percentile were selected for the network structure. A soft-thresholding power (β) of 17 was used to establish scale-free topology within the network structure. A signed hybrid weighted gene co-expression network was constructed, and module eigengenes (MEs) representing the first principal component of each gene module were correlated with sample traits. Hub genes were defined for each module as the top 10% genes with highest degree (sum of connection weights of a gene to all other genes in the module) that have module membership > 0.8.

### Library preparation for whole genome bisulfite sequencing

Whole genome bisulfite sequencing (WGBS) libraries were prepared using Swift’s Accel-NGS Methyl-Seq Kit. The quality of the libraries was checked using an Agilent 2100 Bioanalyzer, and library concentration was measured using a Qubit high sensitivity DNA assay. Individually barcoded libraries were pooled for sequencing. The pool was then sequenced on one lane of a NovaSeq 6000 S4 flow cell at PE150 at the DNA Technologies and Expression Analysis Cores at the UC Davis Genome Center. After sequencing, reads were demultiplexed using the bcl2fastq Illumina software. The number of paired-end reads per sample ranged from 73-109 million (Supplementary Table 7B).

### WGBS read alignment, differential methylation analysis, and pathway analysis

The CpG_Me wrapper pipeline (Laufer et al., 2020, Krueger and Andrews, 2011, Martin, 2011, Ewels et al., 2016) was used to align and analyze the WGBS data. Trim Galore (Martin, 2011) was used to trim the reads. To address potential methylation biases at the 5′ and 3′ ends of reads, 10 bases were trimmed from the 3′ end of reads 1 and 2, and 10 and 20 bases were trimmed from the 5′ end of reads 1 and 2 respectively. Bismark (Krueger and Andrews, 2011) was used to align the reads to the *M. mulatta* genome (Mmul10). Bismark was also used for read deduplication and to generate CpG count matrices. MultiQC (Ewels et al., 2016) was used to evaluate read quality and mapping quality. DMRichR (Laufer et al., 2019, Korthauer et al., 2018, Hansen et al., 2012) was used to identify differentially methylated regions between WS exposed and non-exposed samples. Animal of origin was included as a covariate. The default parameters for DMRichR were used for the differential methylation analysis. Bsseq (Hansen et al., 2012) was used to generate individual smoothed methylation values and methylation heatmaps. IPA (QIAGEN Inc., “https://www.qiagenbioinformatics.com/products/ingenuitypathway-analysis”) was used for pathway enrichment analysis. Pathways with an FDR ≤ 0.05 were considered significantly enriched.

### Similarity scoring analysis

Publicly available count matrices were retrieved from Recount3 and used to generate transcriptomic disease signatures for COPD, IPF, and asthma with DESeq2 (Heymann et al., 2017, Kim et al., 2015, Schafer et al., 2017, Wilks et al., 2021). Genes with an FDR ≤ 0.05 were considered significantly differentially expressed. Similarity scoring was performed between the exposure and disease differentially expressed genes using a previously established Jaccard indexing method (Koval et al., 2022). Briefly, differential expression results were first binarized with significant DEGs assigned a value of 1 and non-significant genes assigned a value of 0. Genes were retained if they were identified as DEGs in at least one exposure condition. Jaccard similarity scores were then computed for each exposure pair using the vegdist function from the vegan R package, with the Jaccard index defined as the intersection over the union of DEGs (Oksanen J., et. al., 2025). This metric captured the extent to which gene expression responses overlapped, irrespective of directionality. Similarity scores ranged from 0 (no overlap) to 1 (complete overlap), and the resulting similarity matrix was used to hierarchically cluster exposure conditions. Cluster number was optimized by evaluating within-cluster sum of squares and silhouette width using the fviz_nbclust function from factoextra, and final clustering was performed with the diana function from the cluster package (Kassambara & Mundt, 2016 and v2.1.1; Maechler et al., 2019).

## Abbreviations

AhR: Aryl Hydrocarbon Receptor
ALI: Air–Liquid Interface
BEAS-2B: Bronchial Epithelial Cell Line (SV40-transformed)
CCL2: C-C Motif Chemokine Ligand 2
COPD: Chronic Obstructive Pulmonary Disease
DEG(s): Differentially Expressed Gene(s)
DMR(s): Differentially Methylated Region(s)
EMT: Epithelial–Mesenchymal Transition
EPA: Environmental Protection Agency
FC: Fold Change
FDR: False Discovery Rate
GO: Gene Ontology
IGF: Insulin-like Growth Factor
IPA: Ingenuity Pathway Analysis
IPF: Idiopathic Pulmonary Fibrosis
JAK: Janus Kinase
ME: Module Eigengene
NF-κB: Nuclear Factor Kappa-light-chain-enhancer of Activated B Cells
NHP: Nonhuman Primate
NGF: Nerve Growth Factor
OEHHA: Office of Environmental Health Hazard Assessment
PHF8: Plant Homeodomain Finger Protein 8
PLA2R1: Phospholipase A₂ Receptor 1
PM₂.₅: Particulate Matter ≤2.5 μm in aerodynamic diameter
PTBP3: Polypyrimidine Tract Binding Protein 3
RNA-seq: RNA Sequencing
ROS: Reactive Oxygen Species
SASP: Senescence-Associated Secretory Phenotype
SMAD3: Mothers Against Decapentaplegic Homolog 3
TGF-β: Transforming Growth Factor Beta
TOM: Topological Overlap Matrix
TP53: Tumor Protein p53
WGCNA: Weighted Gene Co-expression Network Analysis
WS: Wood Smoke
WGBS: Whole-Genome Bisulfite Sequencing

## Declarations

### Ethics approval and consent to participate

Not applicable

### Consent for publication

Not applicable

### Availability of data and materials

WGBS and RNA-seq data will be deposited to GEO upon manuscript acceptance. The code used for analyses will be uploaded to GitHub.

### Authors’ Contributions

HJ and CR conceived the study; SPE extracted DNA and RNA from cells and coordinated AB/PAS analyses; APB performed data analysis of initial RNA-seq and WGBS analysis, correlation between RNA-seq and WGBS, and overlap between DMRs; EH performed WGCNA analysis, integration of RNA-seq with WGBS data, and disease signature analysis; EH drafted and edited the manuscript under the supervision of HJ; APB, SPE, CR and HJ edited the manuscript. All authors reviewed the manuscript.

## Acknowledgements

The sequencing was carried out at the DNA Technologies and Expression Analysis Cores at the UC Davis Genome Center, supported by NIH Shared Instrumentation Grant 1S10OD010786-01.

