## Supplementary Information for "Persistent transcriptomic changes following repeated exposure to wood smoke in non-human primate airway epithelial cells"

### Supplementary Tables

**Supplementary Table 1: Lists of DEGs identified at each exposure condition**. A) DEGs following one day of WS exposure. B) DEGs following five consecutive days of WS exposure. C) DEGs following one week of recovery after repeated WS exposure. Genes were considered significant with an adjusted p-value ≤ 0.05 and an absolute fold change ≥ 1.2. D) Overlapping genes between DEGs following five consecutive days of WS exposure and DEGs following one week of recovery after repeated WS exposure. Genes were considered significant with an adjusted p-value ≤ 0.05 and an absolute fold change ≥ 1.2. E. Demographic information on included animals, including age, sex and weight.

**Supplementary Table 2: IPA results of DEGs from each exposure condition**. A) Pathways enriched among DEGs at one day of exposure. B) Pathways enriched among DEGs at five days of exposure. C) Pathways enriched among DEGs at recovery. D) Pathways enriched among the overlapping 77 DEGs between five-day and recovery exposure groups, including 27 DEGs with concordant directionality.

**Supplementary Table 3**: **DMRs identified following five days of WS exposure**. A) DMRs observed following repeated simulated WS exposure in adult airway epithelial cells differentiated *in vitro*. B) Positional overlaps between DMRs associated with early-life wildfire smoke exposure (*in vivo*) and DMRs induced by repeated WS exposure in adult cells (*in vitro*). DMRs were defined as regions with ≥5% methylation difference and empirical p-value ≤ 0.05. Each entry includes genomic coordinates, direction of methylation change, and nearest annotated gene.

**Supplementary Table 4: IPA results for genes annotated to be nearest to DMRs**.

**Supplementary Table 5: Overlaps between genomic features annotated to DMRs and DEGs.** A) Overlapping DEGs and DMRs annotated to be nearest DEGs following 5 days of WS exposure. B) Overlapping DEGs following 5 days of WS exposure and DMRs annotated to be nearest DEGs following one week of recovery from WS exposure. Overlapping genes are listed with associated fold-change in expression and methylation differences. C) Correlations between DNA methylation changes and expression of nearest genes they were annotated to.

**Supplementary Table 6: IPA results for WGCNA module genes.** A) Genes from the darkorange module are listed alongside associated enriched pathways. B) Genes from the darkorange2 module are listed alongside associated enriched pathways. C) Genes from the lightyellow module are listed alongside associated enriched pathways.

**Supplementary Table 7: Read results for RNA-sequencing and WGBS.** A) The number of paired-end reads per RNA-sequencing sample. B) The number of paired-end reads per WGBS sample.

**Supplementary Figure 1. Combined alcian blue and Periodic Acid-Schiff Staining of ALI cultures following repeated exposures to filtered air and wood smoke.** Representative cultures from each treatment group at 5-day time point and recovery time point are shown in A) and B), respectively.
