## Supplementary figures and images for "Persistent transcriptomic changes following repeated exposure to wood smoke in non-human primate airway epithelial cells"

### Supplementary Figure 1

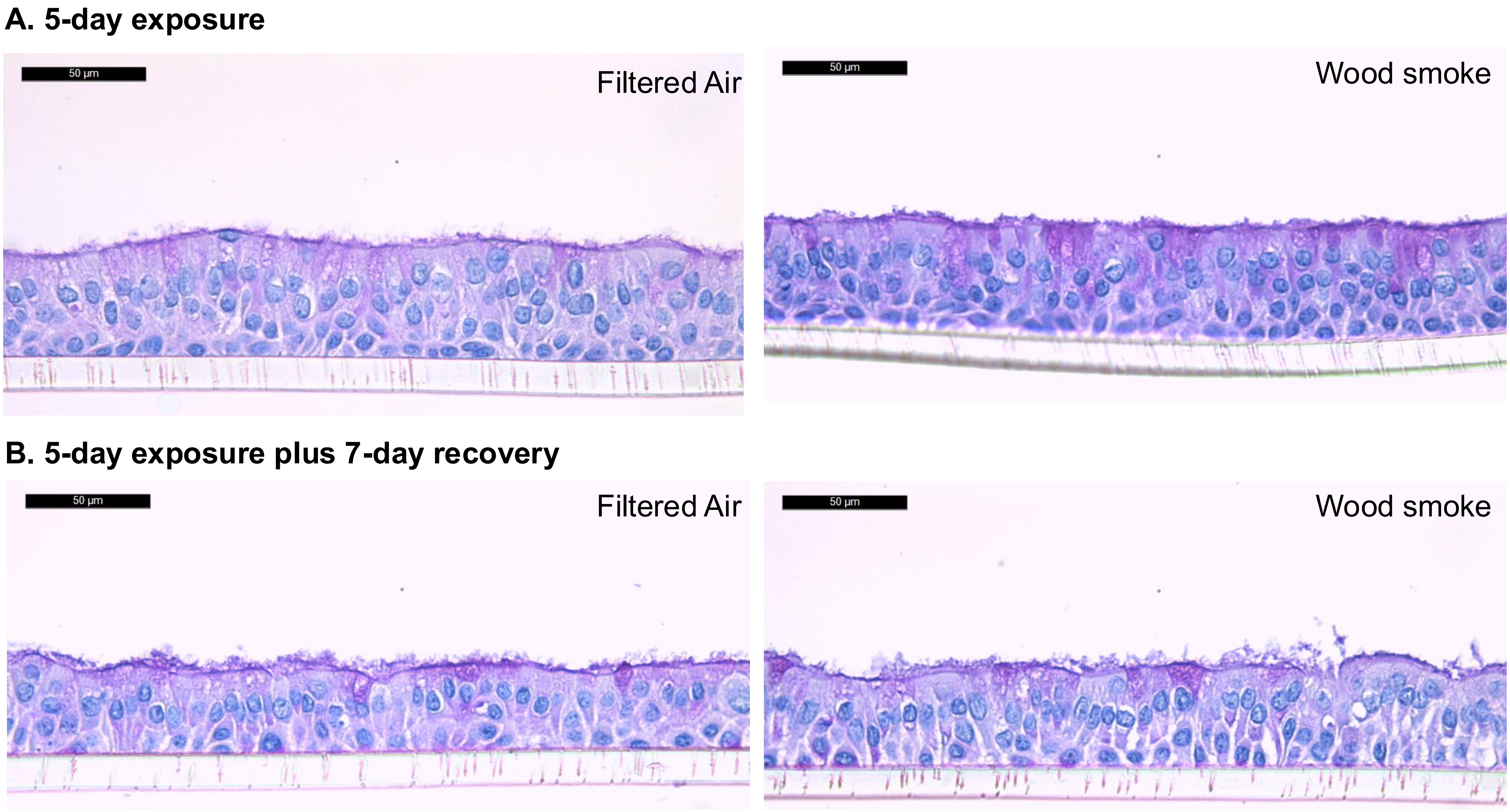
